# Allo-defensive, multiplex base-edited, anti-CD38 CAR T cells for ‘off-the-shelf’ Immunotherapy

**DOI:** 10.1101/2025.08.20.671266

**Authors:** R. Preece, O. Gough, A. Joshi, R. Kadirkamanathan, E. Cudworth, D. Kallon, C. Georgiadis, W. Qasim

## Abstract

Chimeric antigen receptor (CAR) T cell therapies are being widely investigated in both autologous and allogeneic settings, with gene editing providing new strategies to address barriers to mismatched cell therapies. Currently ‘universal’ donor derived T cell therapies require intensive lymphodepletion and are prone to host-mediated rejection. CD38, a transmembrane glycoprotein involved in cell activation and bioenergetics, is a promising immunotherapy target for haematological malignancies. Disruption of CD38 expression using base editing prevented fratricide between T cells expressing anti-CD38 CAR (CAR38). Additional base editing enabled generation of a ‘universal’ donor CAR38-T cells, devoid of endogenous TCRαβ and Human Leukocyte Antigen (HLA) molecules after disruption of *T Cell Receptor Beta Constant* (*TRBC*), *Beta-2 microglobulin* (*B2M*), and *Regulatory Factor X5* (*RFX5*). Removal of cell surface HLA expression enabled evasion of anti-HLA antibodies in sera from sensitised donors and reduced allo-stimulation in mixed lymphocyte cultures (MLCs), while TCRαβ disruption prevented allo-reactivity. In MLCs, CAR38 expression enabled potent ‘allo-defense’ activity against CD38^+^ allo-reactive cells. Multiplex-base-edited CAR38-T cells exhibited antigen-specific anti-leukemic activity against human B, T, and myeloid malignancies and inhibited disease progression in humanised murine xenograft models. CAR38-T cells offer a potent ‘off-the-shelf’ strategy against CD38^+^ haematological malignancies and plasma cells associated with autoimmunity.

## Introduction

Chimeric antigen receptor (CAR) T cells offer new avenues for B cell malignancies including acute lymphoblastic leukaemia (ALL), with products targeting CD19 or B-cell maturation antigen (BCMA) commercially available (1–4). Therapeutic applications are also under investigation for auto-immune disorders where CAR-mediated B cell elimination has induced remissions of Systemic Lupus Erythematosus (SLE) and other conditions (5–8). Limitations and challenges of autologous approaches include disease and host immune cell heterogeneity (9–12), unwanted toxicities from shared antigens (13), and risks of antigen-masking following accidental transduction of blasts (14). Genome-edited allogenic CAR T cells manufactured from healthy donors offer ‘off-the-shelf’ alternatives that can be pre-manufactured and used for a variety of indications (15–18). We have previously manufactured allogeneic anti-CD19 CAR T cells using TALEN or CRISPR/Cas9 editing to remove the T-cell receptor-αβ (TCRαβ) preventing graft-versus-host disease (GVHD) and CD52 to promote survival in the presence of the lymphodepleting antibody Alemtuzumab (15, 16). We have also applied base edited anti-CD7 CAR T cells (BE-CAR7) (13, 17), and BE-CAR33 T cells in human studies against T-ALL and AML respectively. For all these settings, healthy donor derived allogenic CAR (allo-CAR) T cells mediated potent anti-leukemic effects but depended on intense lymphodepletion with augmented doses of fludarabine and cyclophosphamide as well as alemtuzumab. Here we report T cells armed with an anti- CD38 CAR, generated using base editing to first remove CD38 expression, can mediate potent anti-leukemic effects and acquire allo-defensive properties.

CD38 is an extracellular type-II glycoprotein with multiple immune regulatory functions and has been exploited as an immunotherapy target using anti-CD38 monoclonal antibodies daratumumab or isatuximab, which have been approved for indications including multiple myeloma (MM) (19, 20), acute lymphoblastic leukaemia (ALL) and acute myeloid leukaemia (AML) (21–23). Phase-I clinical trials have also reported autologous anti-CD38 CAR (CAR38) T cells with encouraging safety and efficacy profiles (24–28). Genome editing now offers opportunities to improve CAR38 products by disrupting CD38 expression to prevent fratricide and address barriers to allow mismatched allogenic T cells to be used without Human Leukocyte Antigen (HLA) matching.

We combined lentiviral vector delivery of CAR38 with cytosine deaminase mediated base editing to knockout TCRαβ and CD38 alone or in combination with HLA disruption, for universal configurations (***Figure 1A***) (29). Cytidine to thymidine (C>T) conversions introduced premature stop codons or disrupted splice sites of one or more genes at high efficiency and without DNA breaks, allowing BE-CAR38 T cell products to be generated efficiently for investigations *in vitro* and in humanised murine models.

**Figure 1.**
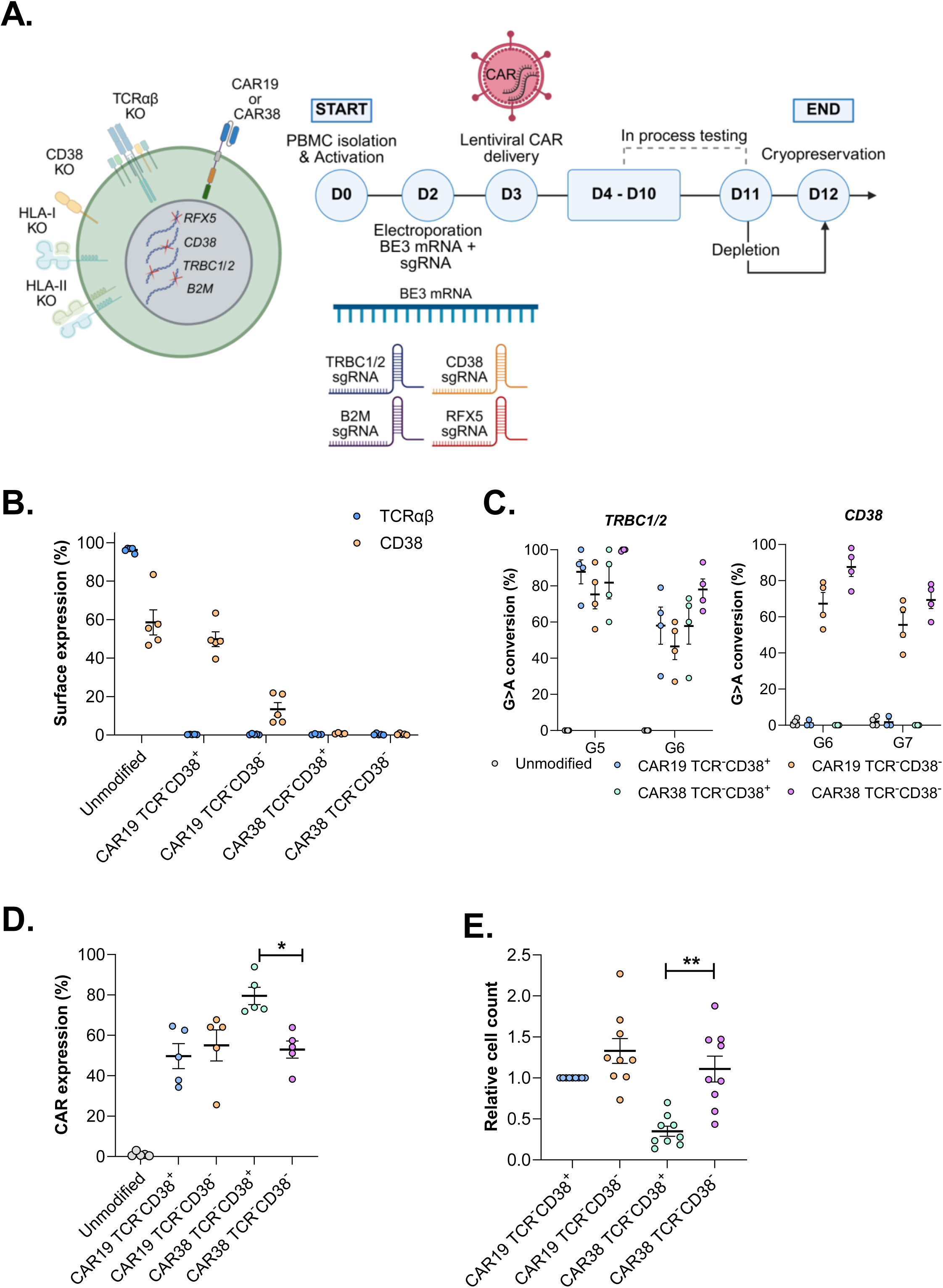
Production of BE-CAR T cells: A) Timeline of base-edited CAR T cell manufacture. B) Quantification of TCRαβ and CD38 disruption at the end of manufacture by flow cytometry (n=5) and C) Sanger sequencing (n=4). D) CAR positive cells post magnetic bead mediated depletion of residual TCRαβ^+^ cells across multiple donors (n=5). * p ≤ 0.05, paired t test. E) End of manufacture cell yields relative to parallel productions of control BE-CAR19 T cells (n=9 donors). ** p ≤ 0.01, paired one-way ANOVA with a Tukey’s multiple comparison test. All data presented as mean ± standard error of the mean.

## Methods

### Cell lines

Cell lines (293T, K562, Daudi, Jurkat, and MOLM14 cell lines) were obtained from ATCC (Virginia, USA). K562, Daudi, Jurkat, and MOLM14 suspension lines were maintained in RPMI-1640 (Thermo Fisher Scientific, Massachusetts, USA), while 293T cells were cultured in DMEM (Thermo Fisher Scientific), both were supplemented with 10% FBS (Sigma-Aldrich, Missouri, USA). Transduction with a lentiviral vector expressing enhanced green fluorescent protein (EGFP) and luciferase (LUC) allowed detection via flow-cytometry and *in vivo* tumour tracking.

Target antigen negative populations (Daudi CD19^-^CD38^-^, Jurkat CD7^-^CD38^-^, and MOLM14 CD33^-^CD38^-^) were generated by CRISPR/Cas9 editing using *Sp*Cas9 mRNA (TriLink BioTechnologies, California, USA) with single-guide RNAs (sgRNA, Synthego, California, USA) specific for CD19, CD7, CD33, or CD38, delivered using a Lonza 4D-Nucleofector X unit (Lonza Group AG, Basel, Switzerland). Residual antigen positive populations were removed by sorting on a MoFlo XDP cell sorter (Beckman Coulter, California, USA).

### Base editing

Codon optimised cytidine base editor-3 (BE3) was described previously (13, 17) and was combined with sgRNAs to create premature stop codons or disrupt splice-donor site as detailed in Supplementary Table S1.

### CAR lentiviral vectors

CARs were expressed under the control of the hPGK promoter in a 3^rd^-generation lentiviral vector configuration (pCCL) and comprised a single-chain variable fragment (scFv) binder region fused to a CD8α derived hinge/transmembrane region, 4-1BB co-stimulatory domain, CD3ζ intracellular signalling domain (scFv-CD8α-4-1BB-CD3ζ) as previously described (13, 17). Anti-CD38 scFv was derived from Daratumumab in a heavy variable – light variable orientation with a GGGGS_3_ linker. Previously described configurations were used for CAR19 (clone: 4g7) (15, 16), CAR7 (clone: 3A1e) (13, 17), and CAR33 (clone: My96) (30).

### CAR T cell manufacture

Mononuclear cells (MNCs) were isolated by Ficoll (Sigma-Aldrich) density gradient from healthy blood donations approved by University College London (REC Reference: 25257.001) or from the Nolan Registry (REC Reference No.: 19/LO/0447) and used to manufacture base edited CAR T cells as described (13). Where indicated, T cells were depleted on day 10 for residual TCRαβ and HLA expressing cells using biotin conjugated mouse anti-human TCRαβ (clone: BW242/412, Miltenyi Biotec), human anti-human HLA-ABC (clone: REA230, Miltenyi Biotec), and human anti-human HLA-DR,DP,DQ (clone: REA332, Miltenyi Biotec) according to manufacturer’s instructions (Miltenyi Biotec). T cells were rested overnight before flow-cytometry based phenotyping and cryopreservation on day 11.

### Flow cytometry

CAR T cell phenotyping by flow-cytometry was carried out using a BD FACSymphony cell analyzer (BD, New Jersey, USA). Primary anti-human antibodies used for immunophenotyping are provide in Supplementary Table S2. CAR19 transduction was assessed using Biotin-SP AffiniPure F(ab’), Fragment Goat, anti-Mouse IgG, F(ab’) Fragment Specific antibody (Jackson Immunoresearch, Cambridge, UK) with secondary staining by Streptavidin (BioLegend, California, USA). His-tagged recombinant CD7, CD33, and CD38 proteins (Sino Biological, Beijing, China) were used to detect CAR7, CAR33, and CAR38 expression, followed by staining with Anti- 6X His tag (clone: AD1.1.10, Abcam, Cambridge, UK). All analysis of flow cytometry was done using FlowJo version 10.10.0.

### Quantification of on-target cytosine base-editing

At the end of CAR T cell manufacture genomic DNA was extracted from 1-2x10^6^ cells using the DNeasy blood and tissue kit (QIAGEN, Hilden, Germany). Primers were designed using Primer-BLAST (https://www.ncbi.nlm.nih.gov/tools/primer-blast/) to amplify 500-1000bp across the protospacer site and manufactured by Thermo Fisher Scientific. Sanger sequencing was performed by Eurofins Genomics (Konstanz, Germany) and analysed using EDITR (http://baseeditr.com/). Primers sequences (5’- 3’) are showed in Supplementary Table S3.

### Flow-based *in vitro* cytotoxicity assay

EGFP expressing antigen positive and antigen negative tumour cell lines were mixed at a 1:1 ratio prior to addition of CAR^+^ effector T cells at a range of effector to target (E:T) ratios. Effectors and targets were co-cultured for 4-hours before staining with a live dead dye (BD) and the target antigen. Cells were then run on the BD FACSymphony cell analyzer (BD), with the ratio of antigen positive to antigen negative tumour cells used to calculate specific lysis.

### Cytometric bead array to quantify cytokine release

Unmodified or CAR T cell groups were cultured alone or co-cultured with target cell lines at an E: T of 1:1 (1x10^5^ each), for 16 hours at 37°C in 200ul RPMI (10% FCS) in a flat-bottom 96-well plate. All conditions were setup in technical triplicate. Supernatant harvest was performed by spinning the plates at 400g for 5 minutes before transferring 150µl to a fresh 96-well plate. Sample preparation and acquisition was performed using the Human Th1/2/17 CBA kit (BD) and a FACSymphony cell analyzer (BD). Data was then analysed using BD FCAP Array software v3.

### Metabolic analysis by Seahorse XF analyser

A XFe96 Sensor Cartridge (Agilent Technologies, California, USA) hydrated with 200µl of calibration fluid (Agilent Technologies) per well and a XFe96 PDL cell plate (Agilent Technologies) were incubated overnight prior to the assay in a 37°C non-CO_2_ incubator. T cells were resuspended at 2x10^6^cell/ml in Seahorse XF RPMI medium supplemented with 1% Seahorse XF Glucose (1.0M solution), 1% Seahorse XF Pyruvate (100mM solution), and 1% Seahorse XF L-Glutamine (200mM solution) all supplied from Agilent Technologies and seeded 1x10^5^ cells per well of XFe96 PDL cell plate (50µl). Assay was performed using the Seahorse XF T Cell Metabolic Profiling Kit (Agilent Technologies) in a XFe96 analyser measuring both extracellular acidification rate (ECAR) and oxygen consumption rate (OCR) in real time. Data was analysed in the online seahorse analytics portal (https:// seahorseanalytics.agilent.com).

### Mixed lymphocyte cultures

Target BE-CAR T cell groups were thawed and rested overnight in TexMACS (Miltenyi Biotec) supplemented with 3% human serum (Seralab) prior to being resuspended at 1x10^6^ cells/ml and irradiated at 30Gy. Irradiated target cells were co-cultured in a U-bottom 96 well plate at a 1:1 ratio (1x10^5^ cells) with freshly isolated PBMCs from an allogenic donor. After 5 days of co-culture, wells are pulsed with 1µCi 3H-thymidine (Revvity, Massachusetts, USA) and incubated for a further 18-20 hours before transfer of 3H-thymidine labelled DNA to a Flitermat (Revvity) using a cell harvester (TOMTEC Imaging Systems, Unterschleissheimn Germany). Meltilex (Revvity) was applied to the Filtermat, and the 3H-thymidine incorporation as a measure of proliferation was read using a MicroBeta counter (PerkinElmer, Massachusetts, USA). For flow cytometry-based readouts co-cultures were setup as above prior to staining and acquisition on day 5 using a BD FACSymphony cell analyzer (BD).

### Natural Killer cell degranulation

Natural Killer (NK) cells were isolated from fresh MNCs using the NK Cell Isolation Kit (Miltenyi Biotec). NK and BE-CAR T cell groups were co-cultured in a U-bottom 96 well plate at a 1:1 ratio (1x10^5^ cells) for 20 hours in TexMACS (Miltenyi Biotec) supplemented with 3% human serum (Seralab). CD107a (clone: H4A3, BioLegend) antibody was added to co-cultures and incubated for an additional 4 hours. Cells were then stained with an antibody panel and acquired on a BD FACSymphony cell analyzer (BD).

### Flow cytometric crossmatch

Five human sera were identified with anti-HLA antibodies collectively covering all HLA molecules present on our BE-CAR T cell donor. Briefly, 4x10^6^/ml BE-CAR T cells were incubated with negative control or test sera for 30 minutes at room temperature. Samples were stained with anti-Human IgG, CD3, and CD19 for 25 minutes in the dark at room temperature. Samples were washed and acquired on a FACSLyric Flow Cytometer (BD).

### *In vivo* CAR T cell studies

Animal studies were approved by the UCL Biological Services Ethical Review Committee and licensed under the Animals (Scientific Procedures PP5675666) Act 1986 (Home Office, London, United Kingdom). NOD/SCID/γc–/– (NSG) mice (Charles River, The Jackson Laboratory), were inoculated by intravenous (IV) injection with either Daudi (CD19^+^CD38^+^, 0.5x10^6^), Jurkat (CD7^+^CD38^+^, 1x10^7^), or MOLM14 (CD33^+^CD38^+^, 1x10^5^) on day 0. All tumour lines were EGFP^+^LUC^+^ and engraftment was confirmed on day 5 by bioluminescence imaging (BLI) using an IVIS Lumina III In vivo Imaging System (PerkinElmer, live image version 4.5.18147). Mice were then injected on day 6 with either 2.5 × 10^6^ unmodified T cells, or 2.5x10^6^ BE-CAR^+^ T cells (up to a maximum of 1×10^7^ total cells). Analysis of tumour inhibition was performed by serial BLI and assessment of bone marrow (processed by a RBC lysis, followed by staining for flow cytometry) at necroscopy.

### Statistical analysis

Means ± standard error of the mean (SEM) are presented and paired or unpaired one-way ANOVA with a post-hoc Tukey’s multiple comparison was performed where indicated. Log rank survival analysis compared outcomes between animal groups. All statistical analysis was performed in GraphPad Prism Version 10.4.1.

## Results

### Base editing prevents fratricide and enables efficient anti-CD38 CAR T cell production

After activation with anti-CD3/CD28 transact reagent T cells upregulated cell surface expression of CD38 from around 11% to 85% (n=3) compared to around 20% (n=3) in T cells with disrupted *CD38* through the introduction of a premature stop codon by base editing (***Supplementary figure 1A&B***). Sanger sequencing confirmed appropriate C>T conversions in the anticipated window of deamination with 77% conversions at position C6 (protospacer position six) and 68% at position C7 (n=3) (***Supplementary figure 1C***). The effects of CD38 knockout were investigated in CAR19 and in CAR38 T cells after transduction with the respective lentiviral vector with simultaneous genome editing of CD38 and TCRαβ/CD3 to create TCRab^-^CD38^-^ CAR^+^ effectors. All BE-CAR groups exhibited high levels of TCRαβ knockout, resulting in ∼1% residual expression after bead mediated depletion. In CAR19 T cells, expression of CD38 reduced from 49.8% to 13.4% (n=5) after *CD38* base-editing, and for BE-CAR38 T cells, there was near complete absence of CD38 expression reflecting ‘self-enrichment’ and/or antigen masking mediated by CAR38 (n=5, ***Figure 1B***). Molecular analysis confirmed flow evidence of editing at *T Cell Receptor Beta Constant 1/2* (*TRBC1/2*) and *CD38* loci with C>T deamination within the anticipated 5bp base-editing window (n=4, ***Figure C***).

Control BE-CAR19 T cells, exhibited similar levels of CAR19 transduction and cell yields irrespective of CD38 base-editing (***Figure 1D & E***) and the phenotype of BE- CAR19 T cells was unaffected by CD38 knockout (***Figure 2A & B***). In contrast, CAR38 T cell yields were significantly greater for TCR^-^CD38^-^ groups compared to TCR^-^CD38^+^ T cells. Although high initial transduction efficiency was documented in the CD38^+^ group, these cells were highly activated (high CD25) and released cytokines (***Figure 2B i***) in the absence of target cells and had an increased bioenergetic profile on Seahorse analysis (***Figure 2C***). These difference in phenotype, activation, and bioenergetic profiles were likely due to CAR38 T cell activation during fratricidal effects against CD38. In contrast TCR^-^CD38^-^ CAR38 T cells released cytokines only in the presence of CD19^+^CD38^+^ Daudi target cells and profiles were comparable to CAR19 controls (n=4) (***Figure 2B ii***). Favourable metabolic profiles have been reported after CD38 disruption in T cells and NK cells (31, 32), however immediately after CAR T cell engineering no significant difference in oxidative phosphorylation (measured by oxygen consumption rate, n=3, ***Figure 2C i***) or glycolysis (measured by extra-cellular acidification rate, n=3, ***Figure 2C ii***) were apparent in BE-CAR19 with or without CD38 knockout. CAR19 and CAR38 effector groups all showed similar lysis of CD19^+^CD38^+^ Daudi target cells across of range of effector: target (E:T) ratios in cytotoxicity assays (***Figure 2D***).

**Figure 2.**
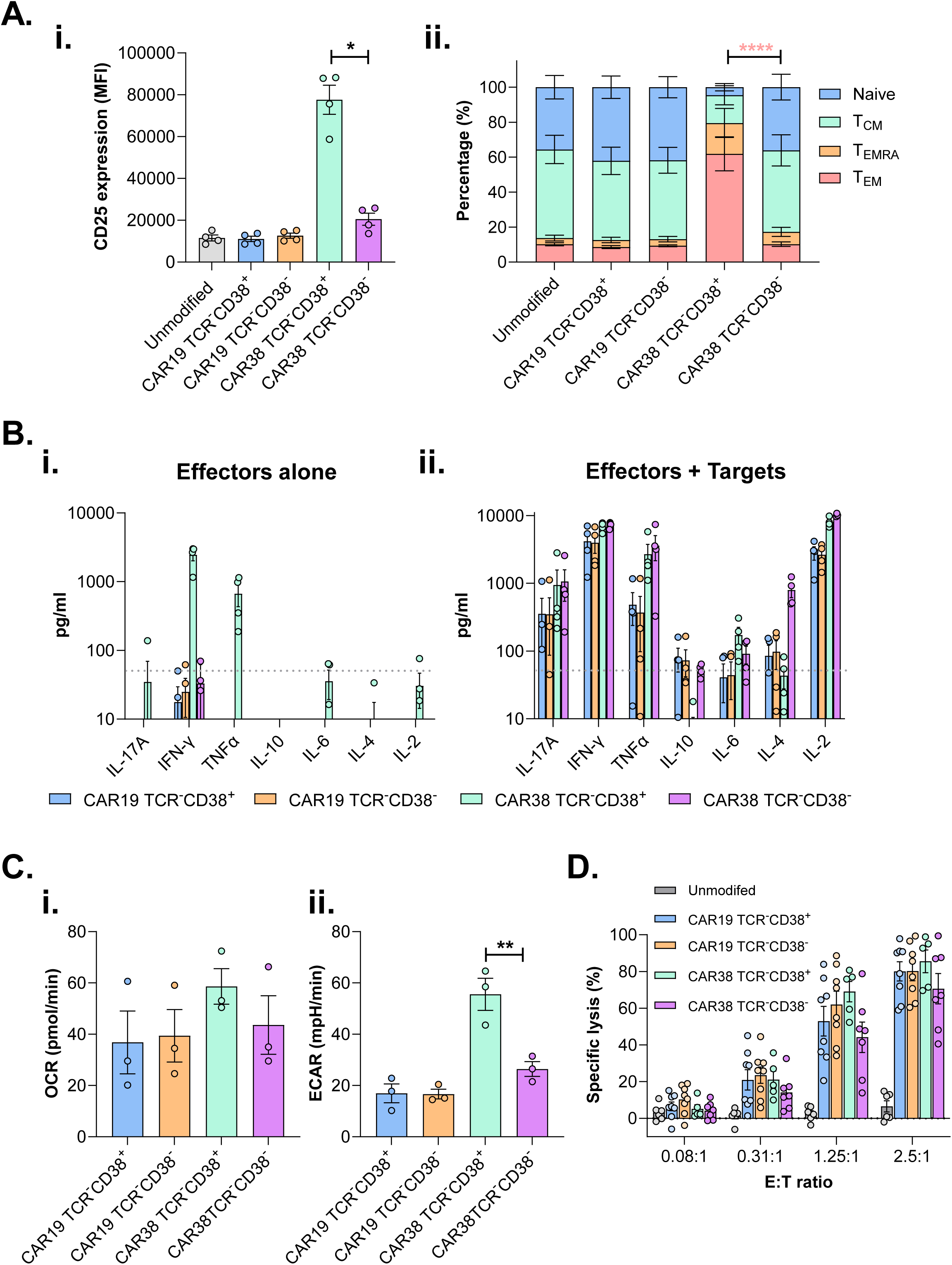
CD38 knockout mitigates against CAR mediated activation and fratricidal effects: A) Immunophenotyping at the end of BE-CAR manufacture (n=4) measuring both activation profile (i, CD25 mean fluorescence intensity; MFI) and memory phenotype (ii, CD45RA and CD62L expression). Naïve (CD62L^+^CD45RA^+^), central memory T cells (T_CM_, CD62L^+^, CD45RA^-^), effector memory T cells re-expressing CD45RA (T_EMRA_, CD62L^-^, CD45RA^+^), and effector memory T cells (T_EM_, CD62L^-^, CD45RA^-^). **** p ≤ 0.0001 paired one-way ANOVA with a Tukey’s multiple comparison test. Red * indicates analysis between the T_EM_ populations. B) Cytokine bead array quantifying cytokine release in absence (i) and presence (ii) of leukaemia target cells (n=3 technical replicates, dotted line indicates the limit of quantification (50pg/ml). C) Metabolic activation measured using Seahorse-XF analyser (n=3 donors, where each point represents the mean of 4-8 technical replicates) measuring oxygen consumption rate (OCR) as a surrogate for oxidative phosphorylation (i) and extracellular acidification rate (ECAR) as a surrogate for glycolysis. ** p ≤ 0.01, paired one-way ANOVA with a Tukey’s multiple comparison test. D) Lysis of Daudi cells after 4-hour *in vitro* co-culture with effector T cells across a range of Effector: Target (E:T) ratios (n=8 donors for all groups except CAR38 TCR^-^CD38^+^ which has n=5 donors). Each point represents the mean of a technical triplicate. All data is plotted as mean ± standard error of the mean.

### HLA base edited BE-CAR38 T cells evade humoral and cellular allo-responses

Overcoming HLA-mismatches to allow allogeneic T cells to be used without matching requires TCRαβ disruption to prevent GvHD, and additional editing such conferring resistance to serotherapy or removal of HLA class-I and -II to tackle host mediated rejection. The latter was achieved by editing *B2M* for disruption of HLA class I expression, and editing of *RFX5*, a major transcriptional factor for HLA class II inhibition. Thus, fully ‘universal’ (TCR^-^ CD38^-^HLAI^-^HLAII^-^) BE-CAR38 and BE-CAR19 T cells were generated by multiplexed disruption of *TRBC1/2, B2M,* and *RFX5 as well as CD38,* followed by bead mediated triple depletion of residual TCRαβ, HLA class I (HLA-I), and class II (HLA-II) expressing T cells (***Figure 3A***). Flow cytometry confirmed knockout and enrichment of highly homogenous TCR^-^HLAI^-^HLAII^-^ CAR T cell products (***Figure 3B i & ii***). Preservation of function of BE-CAR19 and BE-CAR38 T cell products was confirmed *in vitro* after combined TCRαβ and HLA knockouts, with no significant difference in cytotoxicity or cytokine release against CD19^+^CD38^+^ Daudi cells (***Figure 3A & B***).

**Figure 3.**
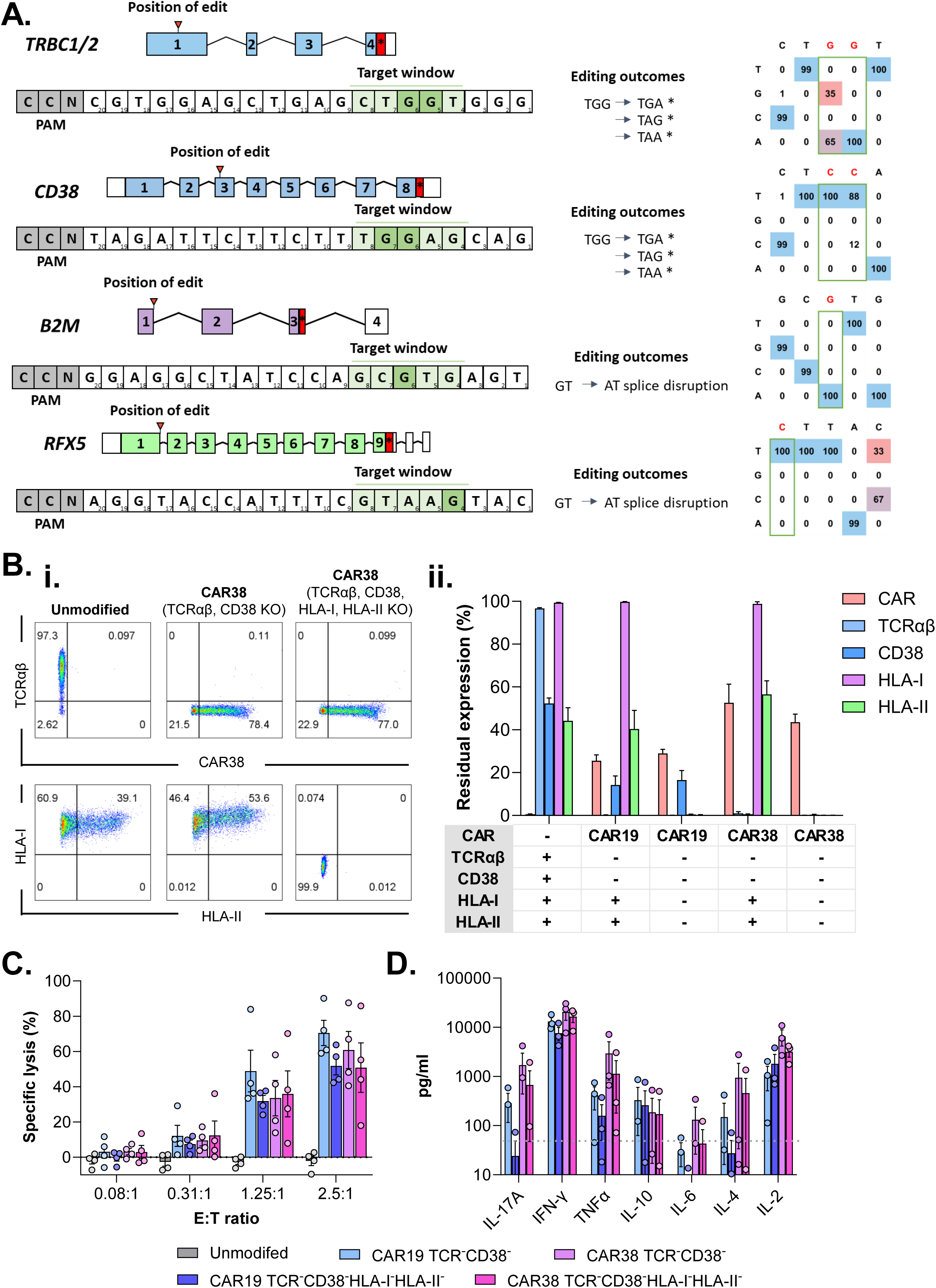
Manufacture of Universal CAR T cells with additional HLA knockouts (TCR^-^CD38^-^HLA-I/II^-^) retain *in vitro* function: A) Schematic of base-edited loci (*TRBC*, *CD38*, *B2M*, & *RFX5*). The red arrow indicates the position of the desired edit, protospacer sequence is shown below, with numbers indicating protospacer position distal to the protospacer adjacent motif (PAM). Line between positions 4 and 8 shows the optimal editing window for the third-generation cytosine base editor (BE3), with targeted bases in dark green. Representative Sanger sequencing analysed by EDITR to quantify base conversion, in CAR38 T cells edited at all loci and depleted for residual TCRαβ and HLA expressing cells. B) Flow-cytometry of unmodified, BE-CAR, and universal CAR T cell groups at the end of manufacture. Representative plots are showing for CAR, TCRαβ, and HLA expression (i) as well as a summary histogram of n=4 donors (ii). C) *In vitro* cytotoxicity of CAR T cell products against a Daudi line in a 4-hour co-culture across a range of Effector: Target (E:T) ratios (n=4 donors, where each point represents the mean of a technical triplicate). D) Cytokine release of effector T cells after 16-hour co-culture with Daudi target cell (n=3 donors). Each point shows the mean of n=3 technical replicates, with the limit of detection indicated by a dotted line (50pg/ml). All data presented as mean ± standard error of the mean.

To investigate cognate recognition and binding of BE-CAR T cells by anti-HLA antibodies we used a flow cross match assay and sera from multiple donors with known HLA sensitisation against the complete repertoire of HLA class I and II molecules expressed by the relevant cell donor (***Figure 4A***). Both BE-CAR19 and BE- CAR38 T cell products with intact HLA expression (BE-CAR19^+^TCR^-^HLAI^+^HLAII^+^ and BE-CAR38^+^TCR^-^HLAI^+^HLAII^+^) exhibited high levels of anti-HLA antibody binding, but in contrast cells with HLA-I and HLA-II knockout exhibited minimal binding that was below limits of quantification. This suggests disruption of both HLA-I and -II offers a route to BE-CAR products even in subjects with pre-existing anti-HLA antibodies.

**Figure 4.**
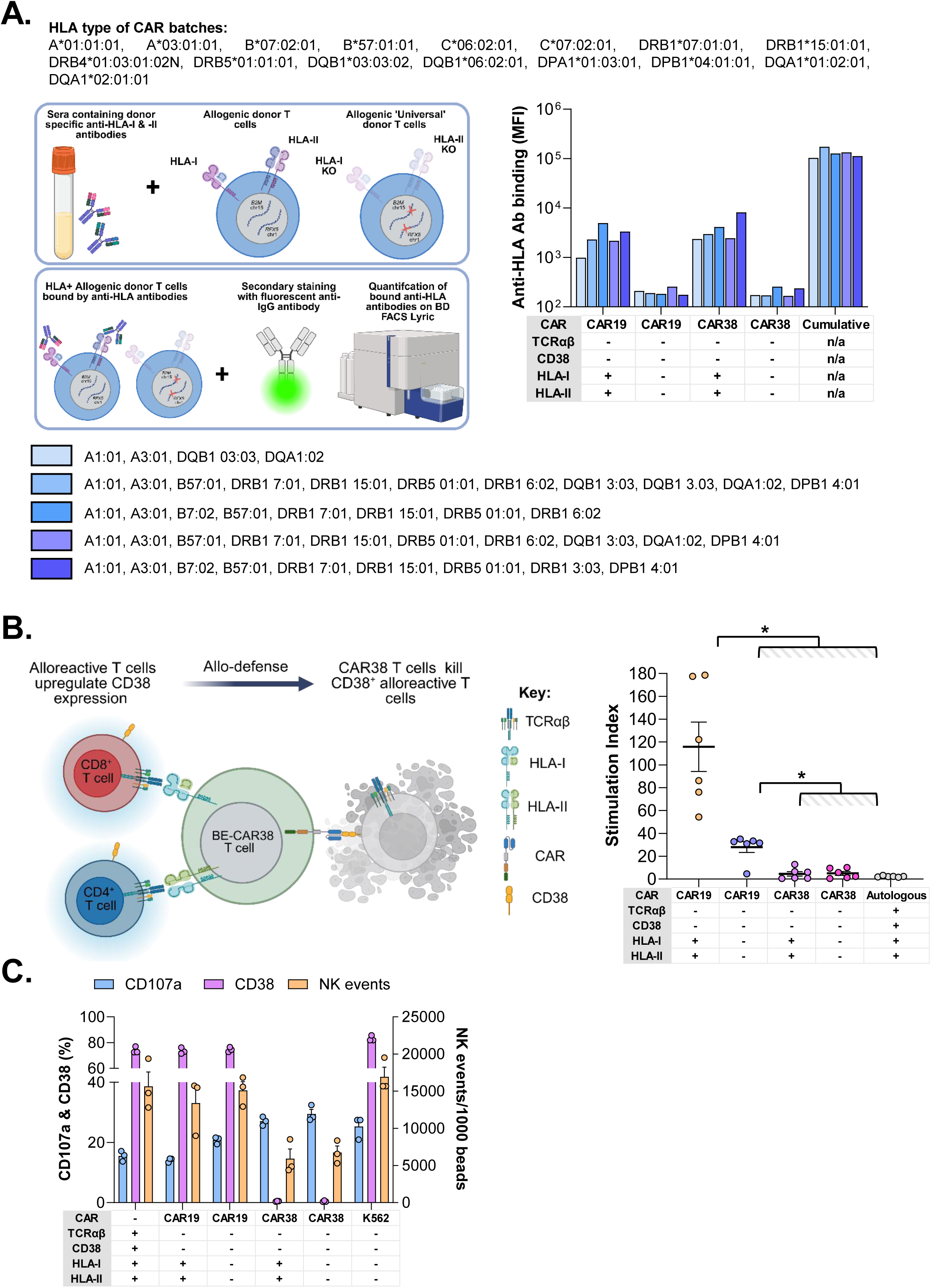
Base-edited TCR^-^CD38^-^HLA-I/II^-^ CAR T cells evade alloreactive humoral and cellular immunity: A) A flow-crossmatch assay was established using human sera with defined anti-HLA antibody profiles. Quantification of anti-HLA antibody binding to BE-CAR T cells from 5 sera measured by mean fluorescence intensity (MFI). B) Mixed lymphocyte reactions measure thymidine uptake by mismatched allo-MNCs against irradiated (30Gy) BE-CAR T cells relative to responses against unmodified T cells (n=6 donors). Each point represents the mean of n=3 technical replicates. One-way ANOVA with a Tukey’s multiple comparison test (*\* P ≤ 0.05*). C) Natural killer (NK) cell degranulation after co-culture with BE-CAR^+^ TCR^-^HLA-I^-^HLAII^-^ T cells or control K562 cells (n=3 donors, with each point representing the mean of a technical triplicate). NK cells are gated on live CellTrace^-^CD2^+^CD4^-^CD8^-^ CD3^-^C56^+^. Expression of CD107a (degranulation marker), CD38 (CAR target), and NK events (measured by counting beads) are plotted for each group. Error bars show mean ± standard error of the mean.

Mixed lymphocyte proliferation assays quantified cell proliferation by 3H-thymidine incorporation as a quantifiable response by host T cell mediated TCRαβ recognition of mismatched HLA on BE-CAR T cells. Thus, these co-cultures assays modelled allo- recognition of irradiated donor BE-CAR T cells by mismatched host T cells and quantified the impact of HLA removal on the allo-stimulation potential of BE-CAR T cells (***Figure 4B***). In the case of BE-CAR19 T cells, disruption of HLA-I and HLA-II significantly reduced responses by non-matched allogeneic T cells. Co-cultures investigating responses elicited against BE-CAR38 T cells revealed more complex interactions involving CAR38 mediated responses against CD38, which was notably upregulated on alloreactive responder T cells. This resulted in significantly reduced thymidine incorporation in proliferation assessments and flow cytometry confirmed that BE-CAR38 T cells eliminated CD38^+^ allogeneic cells in these co-cultures (***Supplementary figure 2***).

Experiments also investigated possible NK cells mediated ‘missing-self’ activity against BE-CAR T cells after HLA-I removal. While there was evidence of NK degranulation against HLA-I^-^ CAR19 T cells, a majority of NK cells expressed CD38 and BE-CAR38 T cells recognised and eliminated these cells in co-cultures (***Figure 4C***). Overall, these data were consistent with CAR38 mediated ‘allo-defense’ phenomena and suggest that ‘universal’ BE-CAR38 T cells may have notable advantages in overcoming host mediated allogeneic responses.

### *In vivo* anti-leukaemia activity of ‘universal’ CD38^-^TCR^-^HLA-I^-^HLAII^-^ CAR38 T cells

To determine *in vivo* antileukemic performance of multiplex edited fully ‘universal’ BE- CAR19 and BE-CAR38 T cells, NSG mice were inoculated with CD19^+^CD38^+^ Daudi cells expressing EGFP and Luciferase six days prior to BE-CAR T cell injections (***Figure 5A*)**. Disease progression was subsequently tracked weekly by IVIS imaging (***Figure 5B***). All groups receiving BE-CAR T cells exhibited significantly reduced disease progression and longer survival compared to mice receiving unmodified T cell controls. Interestingly, survival was longer in CAR38 T cell groups compared to CAR19 groups despite high level expression of both antigens on targets (*P<0.01*, ***Figure 5C & D***). Base editing of CD38 in BE-CAR19 T cells did not appear to influence responses and nor was there significant impact of HLA disruption on function for either CAR product. Flow cytometry of bone marrow at necroscopy detected CAR T cells and quantified residual CD19^+^CD38^+^ Daudi cells (***Figure 5E***), and consistent with BLI, reductions in leukaemia burden were observed to be greatest in the BE-CAR38 T cell group (***Figure 5F***).

**Figure 5.**
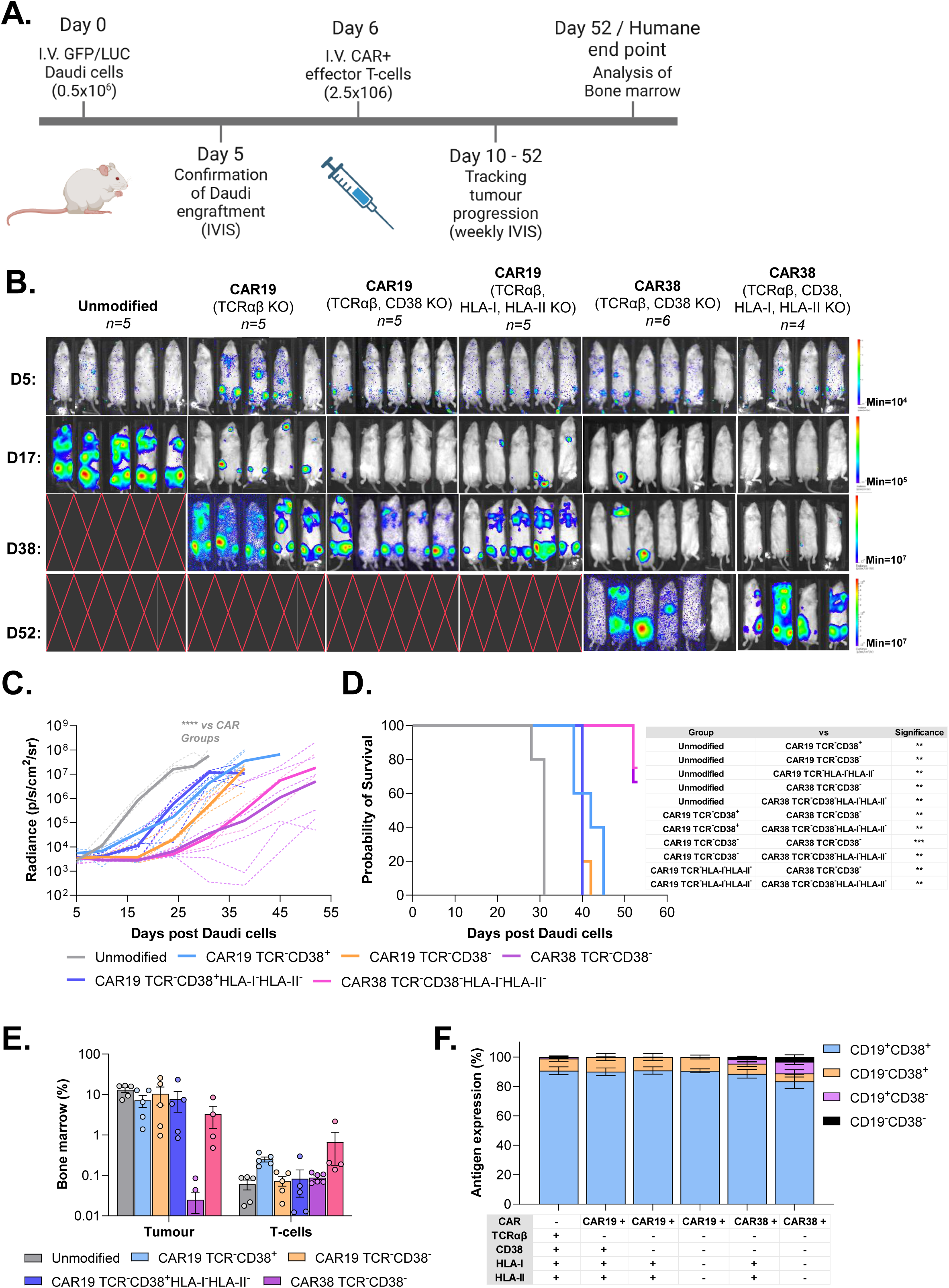
HLA knockout does not impact CAR T cells *In vivo* function: A) Schematic depiction of the *in vivo* experimental timeline with NSG mice engrafted with Daudi cells expressing GFP and Luciferase. B) IVIS images confirming tumour engraftment on day 5, prior to CAR T cell treatment on day 6, and tumour progression over the course of the experiment. Five mice received unmodified and base-edited CAR19 T cells. Six mice received CAR38TCR-CD38-, and four received Universal CAR38 TCR^-^HLA-I^-^II^-^ T cells. C) Leukaemia progression measured by average radiance (p/s/cm^2^/sr) over the course of the experiment. Median of each group is indicated by a solid line, with individual replicates shown as dotted lines. D) Kaplan-Meier curves to day 53, with comparisons between groups performed by log-rank tests. E) Confirmation of both leukaemia burden and residual T cells in the bone marrow at necroscopy. F) Detection of target antigen on remaining GFP^+^ Daudi cells in the bone marrow. Data in E & F is given as mean ± standard error of the mean.

BE-CAR38 T cells were also evaluated against CD7^+^CD38^+^ Jurkat T cell malignant lines and CD33^+^CD38^+^ MOLM14-AML lines, and compared against anti-CD7 CAR T cells (BE-CAR7) (13, 17) and anti-CD33 CAR T cells (BE-CAR33) (30) respectively. *In vitro*, co-cultures with Jurkat (***Supplementary figure 3A***) or MOLM14 (***Supplementary figure 3B***) cells demonstrated antigen specific BE-CAR38 T-cell cytotoxicity and cytokine release against CD38^+^ leukemic lines across E:T ratios. Responses were comparable to BE-CAR7 and BE-CAR33 with no evidence that CD38 editing influenced responses (***Supplementary figure 3A & 3B***). Comparisons *in vivo* used NSG mice engrafted with either Jurkat or MOLM14 lines and again responses were comparable to BE-CAR7 (***Supplementary figure 4A***) or BE-CAR33 respectively (***Supplementary figure 4B***). These findings suggest that BE-CAR38 T cells have potential applicability against a wide variety of CD38^+^ haematological malignancies.

## Discussion

CD38 is a promising candidate for immunotherapy with robust expression in multiple haematological malignancies (21, 22, 33–36) but limited expression on healthy tissue beyond the hematopoietic system (37). Anti-CD38 monoclonal antibodies exemplified by Daratumumab and Isatuximab, received FDA approval for treatment of MM due to their efficacy and safety profiles (38–42), and have also produced encouraging outcomes against ALL (43–45), although resistance and limited clinical responses have also been documented (46). CD38 is also expressed on long lived plasma cells and there is also interest in targeting these populations for certain autoimmune conditions (47).

Alternative CAR T cell based approaches have also been explored with early clinical trials reporting efficacy and safety data in B-ALL (24), AML (25), chronic myelogenous leukaemia (CML) (27), and MM (26, 28). These autologous CAR38 T cell products were reported to have negligible residual CD38^+^ cells detectable by flow cytometry at the end of manufacture, suggesting possible epitope masking by the CAR and/or enrichment of the CD38 negative T cell populations during manufacture (24).

We found that CD38 disruption by base editing improved CAR38 T cell yields significantly by protecting against cell loss through fratricide and reducing metabolic activation, cytokine release and terminal differentiation of effector cells. CD38 enzymatic activity is known to deplete NAD^+^ availability while producing adenosine, which in turn has been associated with T cell immunosuppression (48, 49). Some reports have suggested that inhibition or knockout of CD38 favours oxidative metabolism and offers improved function of CAR T cells (31, 32). We found no evidence that CD38 knockout affected function either *in vitro* or *in vivo* for BE-CAR38 or other CD38 edited CAR T cells targeting CD19, CD7 or CD33. Other strategies for CAR38 T cell or NK-cell approaches have investigated restriction of CD38 with blocking antibody (50), CRISPR/Cas9 genome editing (51–53). affinity-optimised scFvs (54) and adaptor-based CARs (33). We investigated ‘universal’ allogenic CAR38 T cell approach as an ‘off-the-shelf’ alternative that could be applied across a variety of indications. Previously, for universal CAR T cells we combined TCRαβ knockout to avoid GVHD and CD52 knockout to confer resistance to the anti-CD52 monoclonal antibody alemtuzumab, which is used to lymphodeplete recipients and reduce the risk of host mediated CAR T cell rejection. In the context of CD38 knockout T cells it may be feasible to use Daratumumab or Isatuximab instead of alemtuzumab to create a similar advantage for allo-CAR T cells. Moreover, removal of both HLA class I and II was incorporated to extend immunological stealth, allowing evasion of pre-existing anti-HLA antibodies and reducing allo-stimulation likely to trigger host immune cell mediated rejection Multiplexed based editing of *CD38*, *TRBC1/2*, *B2M* and *RFX5* enabled BE-CAR T cells to evade binding by anti-HLA antibodies and ameliorated cell mediated responses in mixed lymphocyte cultures where alloreactive T-lymphocyte recognition of de-nuded BE-CAR19 iterations was blunted. In the context of BE-CAR38 T cells these cells exhibited responses against activated CD38^+^ allogeneic T cells and NK cells, a phenomena akin to ‘allo-defense’. The ‘allo-defense’ concept has been previously described, for example by targeting upregulated 4-1BB on activated lymphocytes, or through B2M - CD3ζ fusion receptor for depleting allo-reactive T cells upon HLA-I recognition (55, 56). In combination, expression of CAR38 and quadruple base-editing has the potential to both arm T cells against and shelter T cells from host immunity. Therapeutic development against a variety of haematological malignancies and autoimmune conditions using ‘universal’ ‘off-the-shelf’ BE-CAR38 T cells are warranted, either alone or in combination with other CD38 edited ‘off-the-shelf’ CAR T products.

## Acknowledgements

All research at Great Ormond Street Hospital NHS Foundation Trust and UCL Great Ormond Street Institute of Child Health is made possible by the NIHR Great Ormond Street Hospital Biomedical Research Centre. The views expressed are those of the author(s) and not necessarily those of the NHS, the NIHR or the Department of Health. This work has been supported by Wellcome Trust (215619/Z/19/Z). Neuza Pina at the Clinical Transplantation Laboratory, Barts Health NHS Trust assisted in running the flow cytometric crossmatch assay. Hannah Rosa and Aakruti Kaikini at King’s College London helped with the Seahorse XF analyser. Ayad Eddaoudi and Panayiota Constantinou, in the Flow Cytometry Core Facility at UCL GOS Institute of Child Health, supported by the Great Ormond Street Children’s Charity (GOSHCC) provided flow cytometry support. Ailsa Greppi and Kyle O’Sullivan supported *in vivo* studies. Anthony Nolan Trust supplied MNC donations.

## Author contributions

W.Q., R.P., and C.G., designed the project. R.P., C.G., O.G., R.K., A.J., E.C., and D.K., performed experiments and analysed data; R.P., and W.Q. wrote the manuscript. All authors have reviewed and approved the manuscript.

## Data availability

Raw data presented and analysed in this manuscript can be requested from the corresponding author.

