## Supplemental Tables and Figure Legends for "Allo-defensive, multiplex base-edited, anti-CD38 CAR T cells for ‘off-the-shelf’ Immunotherapy"

| sgRNA ID | Protospacer sequence (5'-3') | Intended effect |
|---|---|---|
| **TRBC1/2** | CCCACCAGCTCAGCTCCACG | Premature stop codon |
| **B2M** | ACTCACGCTGGATAGCCTCC | Removes splice donor |
| **RFX5** | GTACTTACGAAATGGTACCT | Removes splice donor |
| **CD38 sgRNA 1** | GCGCCAGCAGTGGAGCGGTC | Premature stop codon |
| **CD38 sgRNA 2** | CTCCACTGCTGGCGCCACCT | Premature stop codon |
| **CD38 sgRNA 3** | CTGCTCCAAAGAAGAATCTA | Premature stop codon |
| **CD38 sgRNA 4** | GTTTTCCAGAATACTGAAAC | Premature stop codon |
| **CD38 sgRNA 5** | AGGTTCAGACACTAGAGGCC | Premature stop codon |
| **CD38 sgRNA 6** | TTACCTGTAGATATTCTTGC | Removes splice donor |
| **CD7** | CACCTGCCAGGCCATCACGG | *Sp*Cas9 indel formation |
| **CD19** | GGAACCTCTAGTGGTGAAGG | *Sp*Cas9 indel formation |
| **CD33** | TGACAACCAGGAGAAGATCG | *Sp*Cas9 indel formation |

*Supplementary Table 1: Protospacer sequences used throughout this manuscript compatible with BE3 or SpCas9 genome editing. Sequences are given in 5'-3' orientation, with the target Cytosine highlighted in Red.*

| Anti-human antibody | Clone | Supplier |
|---|---|---|
| **CD2** | LT2 | Miltenyi |
| **CD4** | OKT4 | BioLegend |
| **CD8** | SK1 | BioLegend |
| **CD7** | CD7-6B7 | BioLegend |
| **CD16** | 3G8 | BioLegend |
| **CD19** | HIB19 | BioLegend |
| **CD25** | M-A251 | BioLegend |
| **CD33** | WM53 | BioLegend |
| **CD38** | HIT2 | BioLegend |
| **CD45** | REA747 | Miltenyi |
| **CD45RA** | HI100 | BioLegend |
| **CD56** | REA196 | Miltenyi |
| **CD62L** | REA615 | Miltenyi |
| **CD71** | CY1G4 | BioLegend |
| **CD107a** | H4A3 | BioLegend |
| **HLA-A2** | BB7.2 | BioLegend |
| **HLA-ABC** | W6/32 | BioLegend |
| **HLA-DR, DP, DQ** | Tü39 | BioLegend |
| **TCRαβ** | IP26 | BioLegend |

*Supplementary Table 2: Flow cytometry antibodies used for immunophenotyping.*

| Target locus | Forward Primer (5’-3’) | Reverse Primer (5’-3’) |
|---|---|---|
| **TRBC1/2** | ACACAGAGCCCCTACCAG | GCTACCTGGATCTTTCCA |
| **B2M** | CCTCCAGCCTGAAGTCCTAG | GACGAAGTCCACAGCTCTCC |
| **RFX5** | GTATGGGGTCAGAGGCAGAA | GGGCTTCTATGCAAGTGCTC |
| **CD7** | ATCACCTGCTCCACCAGCGG | GTGTCCTCGCCAGCACACAC |
| **CD33** | CTGTAGTCCTTCCCCTCCAC | CAGCGAACTTCACCTGACAG |
| **CD38** | GGGAGTTAGCGGAGGGAGTA | GCGGAAACCGCAGAAAAAGT |

*Supplementary Table 3: Primer sequences used for genomic analysis of base conversion*

### Figure legends

**Supplementary figure 1 CD38 kinetics on primary human T cells post activation and knockout by base-editing:** A) Flow-cytometry monitoring expression of CD38 and CD25 on T cells ($CD2^{+}CD45^{+}$) from n=3 MNC donors pre- and post-activation up to day 12. B) Schematic of CD38 locus. Exons are shown as solid squares, with joining black lines indicating intronic sequence. Triangles show sgRNA binding sites and predicted outcome of premature stop codon creation or splice donor site (SD) disruption. Flow cytometry measuring residual CD38 expression after cytidine deaminase base-editing across all tested sgRNA sequences (n=3 donors) with sgRNA 3 having the lowest residual expression. C) Sanger sequencing across the protospacer of sgRNA 3 confirming C>T conversion at protospacer positions 6 and 7 within the predicted editing window consistent with flow-cytometry (n=3 biological replicates). Analysis performed using EDITR. All plots are presented as mean ± standard error of the mean.

**Supplementary figure 2 Based-edited CAR38 T cells depleted $CD38^{+}$ allo-responsive T cells in mixed lymphocyte co-cultures:** A) Representative flow-cytometry after 5-day mixed lymphocyte co-culture with responder MNCs and irradicated target cells (30Gy, Unmodified T cells). Responder T cells are gated on live $HLA\text{-}A2^{+}CD2^{+}CD45^{+}TCR\alpha\beta^{+}$, with CD25 and CD71 expression used as measures of alloreactivity. These co-cultures were setup in technical triplicate with responders alone and autologous MNCs as negative controls, allogenic MNCs targets used as positive controls, and BE-CAR groups as test conditions. B) This was performed for n=3 biological replicates. Each point represents the mean of a technical triplicate. Data is plotted as mean ± standard error of the mean.

**Supplementary figure 3 *In vitro* cytotoxic function of BE-CAR38 T cells against Jurkat and MOLM14 lines:** Tumour lysis and cytokine release of unmodified and BE-CAR38 effector T cells (n=4 donors) against A) Jurkat or B) MOLM14 cell line with BE-CAR7 or BE-CAR33 used as a comparison. Each point represents the mean of a technical triplicate. All plots display mean ± standard error of the mean.

**Supplementary figure 4 *In vivo* function of BE-CAR38 T cells in other models of haemopoietic malignancy:** NSG mice were engrafted with either Jurkat (A) or MOLM14 (B) tumour lines ahead of BE-CAR38 effector T cells. Unmodified T cells were given as a negative control, with BE-CAR7 or BE-CAR33 used as a control with known potency. Leukaemia progression measured by average radiance (p/s/cm$^2$/sr) over the course of the experiment. Solid lines present the median of each group with each replicate shown as dotted lines. Survival in each group is shown on Kaplan-Meier curves before reaching a humane end point or the end of experiment. Statistical analysis of survival curves performed by log-rank test.
