## Supplementary figures and images for "Allo-defensive, multiplex base-edited, anti-CD38 CAR T cells for ‘off-the-shelf’ Immunotherapy"

### Supplemental Figure 1

#### Supplementary figure 1:

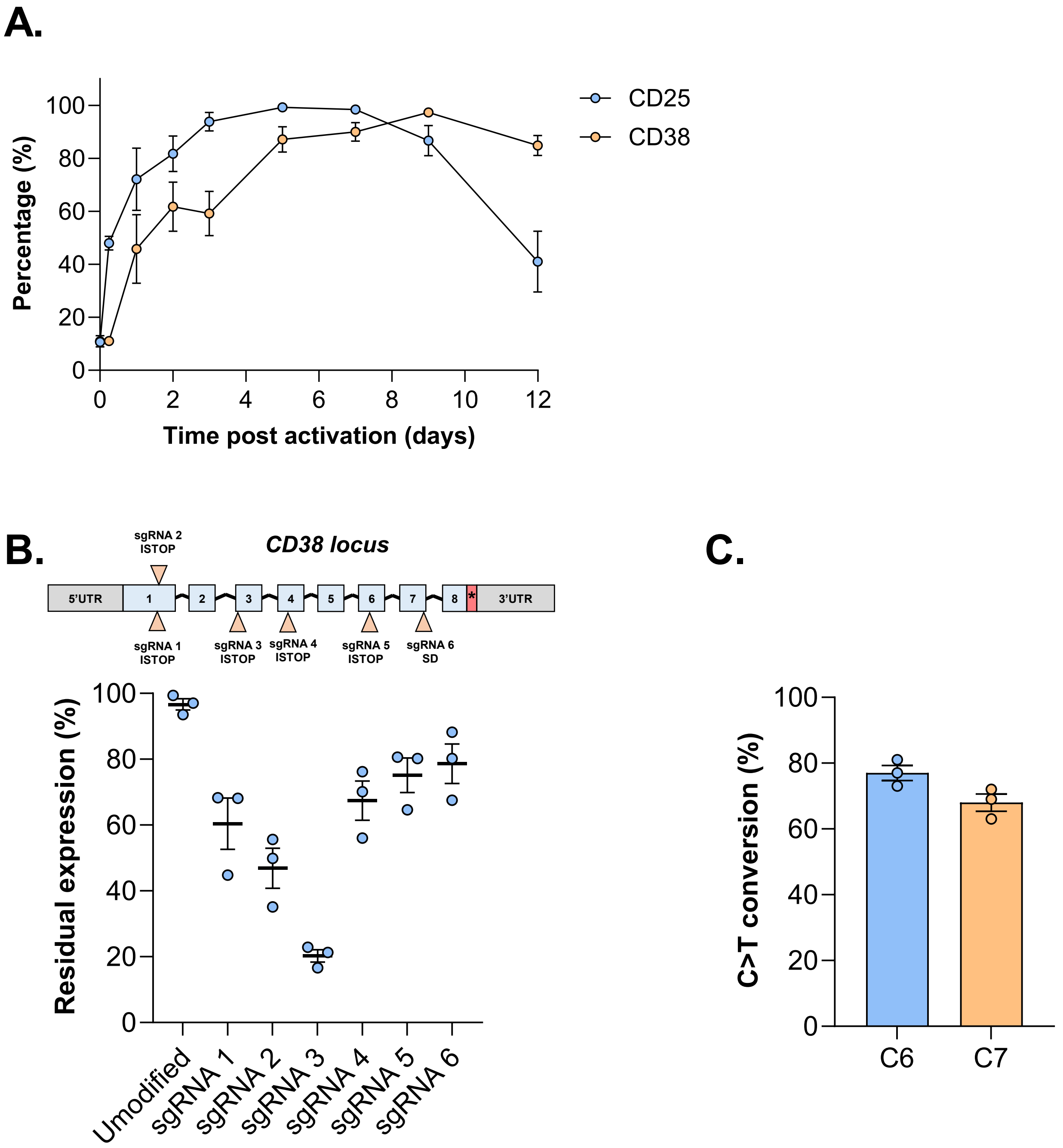

### Supplemental Figure 2

#### Supplementary figure 2:

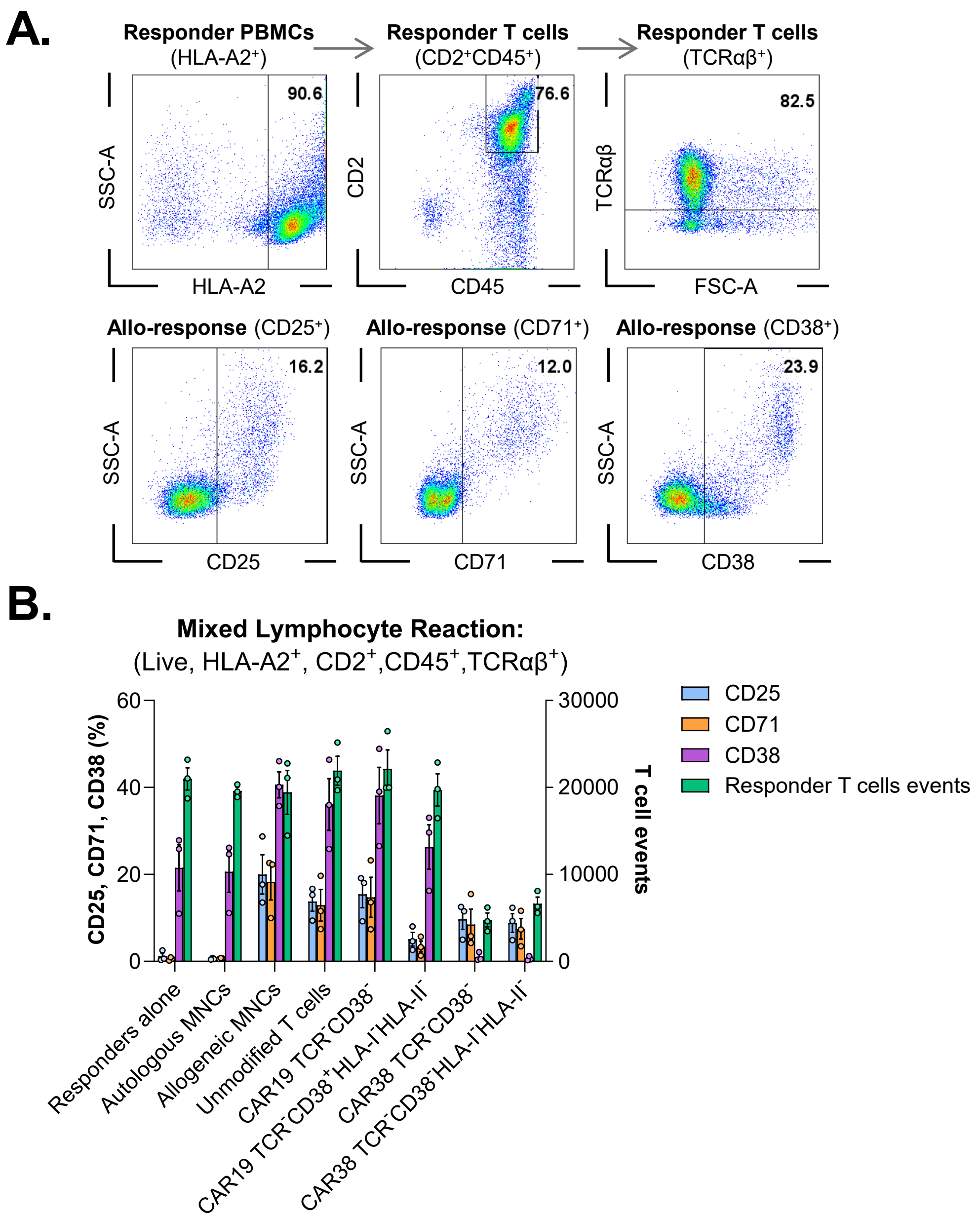

### Supplemental Figure 3

#### Supplementary figure 3:

## A.

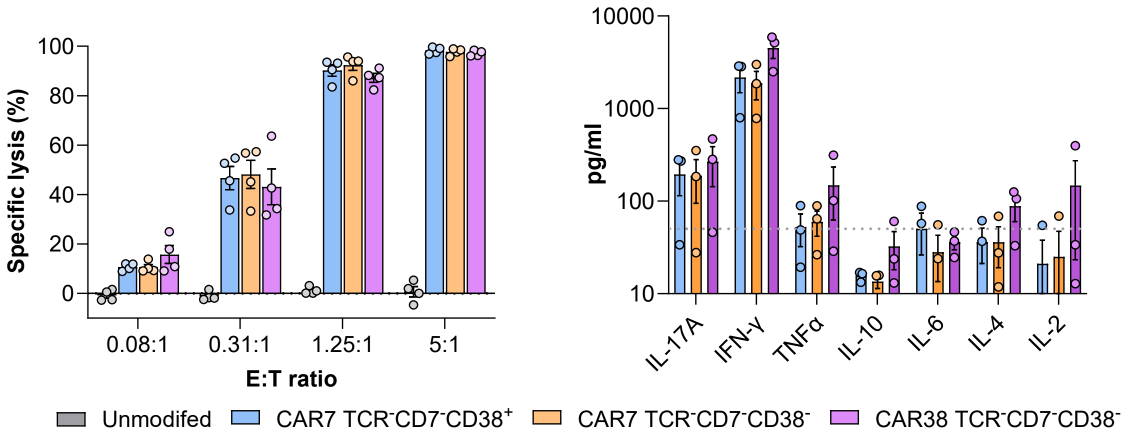


## B.

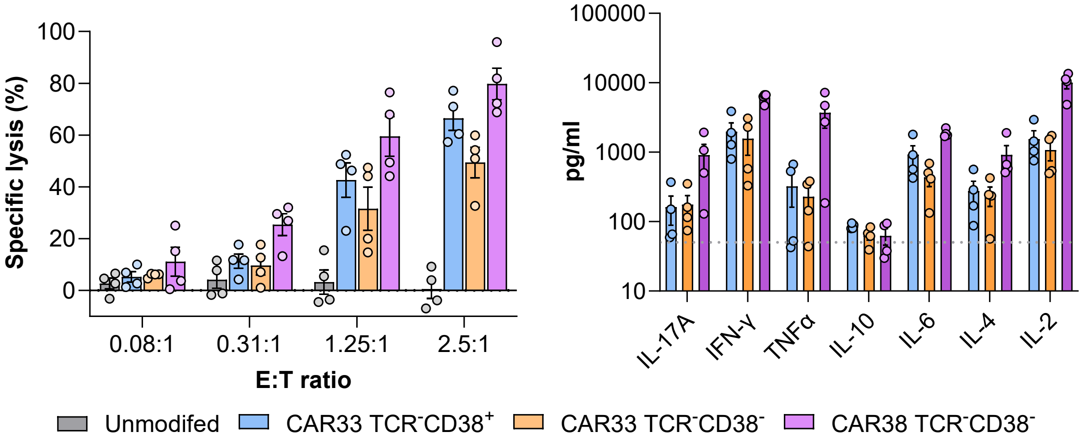
