## Supplemental Figure 4 for "Allo-defensive, multiplex base-edited, anti-CD38 CAR T cells for ‘off-the-shelf’ Immunotherapy"

### Supplementary figure 4:

**A.**

**i.**

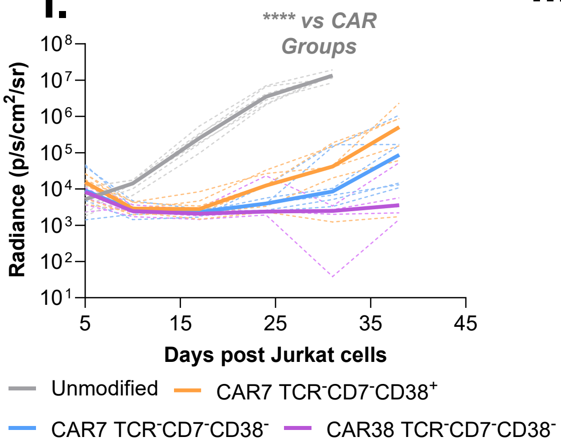


**ii.**

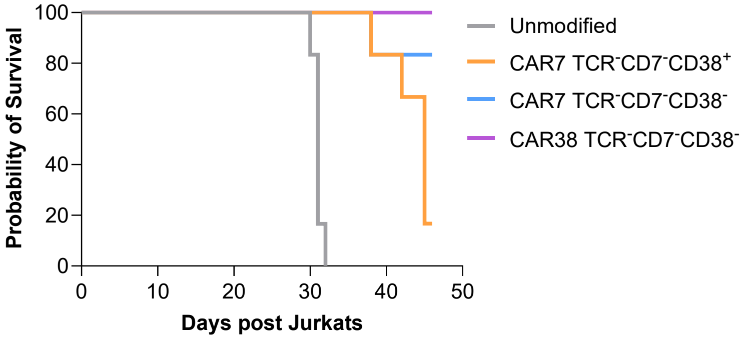


| Group | *Vs* | Significance |
|---|---|---|
| Unmodified | CAR7 TCR$^-$CD7$^-$CD38$^+$ | *** |
| Unmodified | CAR7 TCR$^-$CD7$^-$CD38$^-$ | *** |
| Unmodified | CAR38 TCR$^-$CD7$^-$CD38$^-$ | *** |
| CAR7 TCR-CD7-CD38+ | CAR38 TCR$^-$CD7$^-$CD38$^-$ | ** |

**B.**

**i.**

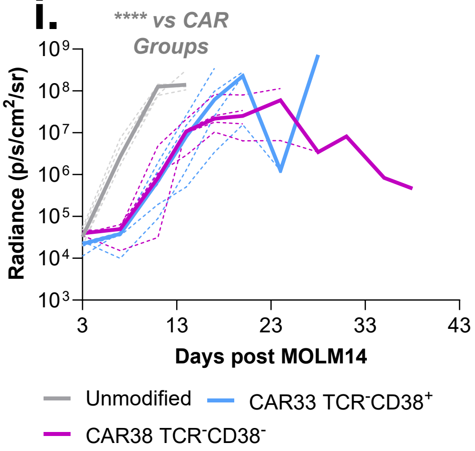


**ii.**

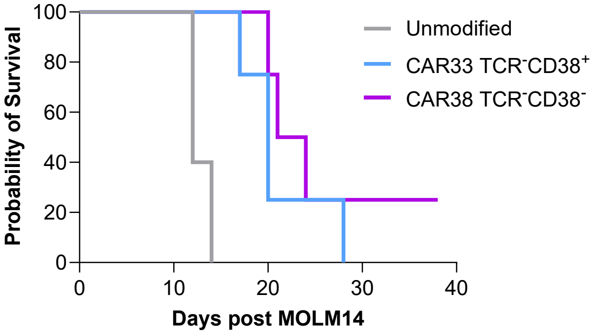


| Group | *Vs* | Significance |
|---|---|---|
| Unmodified | CAR33 TCR$^-$CD38$^+$ | ** |
| Unmodified | CAR38 TCR$^-$CD38$^-$ | ** |
